# Bulked segregant RNA-Seq reveals differential genes and non-synonymous genetic variants linked to sucrose accumulation in sugarcane

**DOI:** 10.1101/792218

**Authors:** Nandita Banerjee, Sanjeev Kumar, A. Annadurai, P. K Singh, J. Singh, R.K. Singh, Sanjeev Kumar

## Abstract

Sucrose is the key economic trait in sugarcane which is a highly polyploid multi-species hybrid and shows a complex pattern of trait inheritance. The excessively large genome of sugarcane is comparatively less explored through Next Generation Sequencing tools for creating superior varieties. In this study, RNAseq libraries of two extreme bulks from a segregating full-sib population and its parents were used to identify 9905 condition-specific non-synonymous genetic variants (NSVs), out of which 43 had a very high degree of differential enrichment (Δf >0.5) for contrasting sucrose accumulating conditions. The statistical model used in this study which was able to quantify the relative effect of NSVs on the trait detected highly significant positive and negative effect NSVs located in the coding regions of genes involved in sucrose metabolism, photosynthesis, mitochondrial electron transport, glycolysis and transcription. In addition, a few differential pre-mature stop codons that could result in production of truncated proteins were also detected in genes coding for aquaporin, GAPDH, aldolase, cytochrome C-oxidase, chlorophyll synthase and plant plasma membrane intrinsic proteins. Additionally, a total of 2140 differentially expressed genes (DEGs) linked to high sucrose accumulation were identified. Among the DEGs, sucrose phosphate synthase III, genes involved in transport, auxin signal transduction, etc., were upregulated, while those involved in electron transfer, cytochrome P450, etc., were downregulated in high sucrose accumulation conditions. This study was able to give finer insights in to the role of allelic heterozygosity on sucrose accumulation and the identified NSVs and DEGs could be useful as candidate markers in marker-assisted breeding for developing high sugar varieties.

## Introduction

Sugarcane (*Saccharum* spp. hybrid) is a crop of unquestionable importance around the globe due to its commercial products like sugar and ethanol, and a number of by-products. It possess one of the most complex and excessively large (map length ∼17000 cM) genomes that makes it difficult to identify genes/causal variants that result in desirable trait expression. Among the sugarcane accessions, the chromosome number vary from 100 to 130 (Butterfield et al., 2001; D’Hont and Glaszmann, 2001), and dozens of copies for each homeologous chromosome pose a formidable challenge in application of Next Generation Sequencing tools. In sugarcane where a reference genome is lacking, pooled RNA-Seq is considered to be the best approach to get a reduced representation of the genotype (Schllotterer et al. 2014). Recent advancements in techniques of SNP detection in the transcriptome of polyploid species provide an opportunity to use NGS-enabled Genetics for identifying causal variants linked to traits of interest (Trick et al. 2012).

Despite the importance of sugarcane, its whole genome sequencing is in the initial phase with the report of a BAC based monoploid genome sequence of sugarcane (*Saccharum spp*. hybrid) that has been aligned on the gene rich part of the sorghum genome (Garsmeur et al. 2018). Recently, an allele-defined reference genome has also been published for *S. spontaneum* (Zhang et al. 2018). Schneeberger and Weigel (2011) suggested that even in absence of complete genome sequence, it might be possible to identify genes underlying traits of interest using bulked segregant RNA seq (BSR-seq) approach. Deep sequencing of transcriptome (RNA-seq) has the potential to uncover the causal variations of a desirable trait both at the genic as well as at allelic level. At the genic level, there have been a few reports in sugarcane on differentially expressed genes (DEGs) associated with to sucrose accumulation (De Setta et al. 2014; Cardoso-Silva et al, 2014; Thirugnanasambandam et al. 2017, 2019), lignin accumulation (Vicentini et al. 2015; Kasirajan et al. 2018), photosynthesis (Mattiello et al. 2015), response to infections from *Gluconacetobacter* (Vargas et al. 2014), and smut pathogen (Wu et al. 2013; Que et al. 2014; Su et al, 2015; Brigida et al. 2016). Recently, Singh et al. (2018) investigated the allele specific expressions linked to molecular mechanism behind high biomass accumulation using BSR-Seq of parents, the F_1_ and 20 F_2_ clones of sugarcane. The type and degree of trait expression is determined by both the structure as well as combination of the genetic elements within the individual, and hence additional studies that could predict and quantify the functional effects of favourable alleles to the expression of a desirable trait could provide deeper insights in to gene regulation. Non-synonymous genetic variant (NSV) is a single nucleotide polymorphism (SNP) that falls within the coding region of a gene and is able to cause an amino-acid change in the protein sequence. In this study, BSRseq was employed to identify differentially expressed genes (DEGs) and NSVs associated with sucrose accumulation using high and low sugar parents and two bulks with extremes for sucrose accumulation obtained from their F_1_ segregating population. The NSVs identified in this study may be used directly as quantitative trait nucleotides (QTNs) that can be translated in to markers for use in marker-assisted breeding program. The results would be useful in future genome assembly and fine mapping in the complex genome of sugarcane and also advance our understanding about interplay of genes involved in early sucrose accumulation.

## Materials and methods

### Plant materials and phenotyping

This study utilized a mapping population comprising of 262 F_1_ individual genotypes developed from a cross between MS 68/47 (low sucrose) × CoV 92102 (high sucrose). The cross was made at ICAR-Sugarcane Breeding Institute, Coimbatore, India in Dec. 2014, while F_1_ plants were grown at ICAR-Indian Institute of Sugarcane Research, Lucknow, India in June 2015. In the first year, both the parental lines along with 262 F_1_ plants were planted at 100 × 90 cm spacing in an augmented block design II. The 262 individuals were planted in four blocks containing 66 individuals in the first three and 64 individuals in the fourth block along with the four check varieties (CoJ 64, BO 91, Co 1148, CoS 767) in each block and recommended agricultural practices were followed to raise a good crop. The true F_1_ seedlings were detected using male parent (MS 68/47) specific 5 EST-SSR markers (Supplementary Table 1).

The mapping population was phenotyped for two consecutive crop cycles (2015-16 and 2016-17). Sucrose content was recorded at 10 month by Nelson-Somogyi method (Somogyi 1952). Corrected Brix, sucrose content (%) and purity in cane (%) was measured for each of the F_1_ plants. Based on the data on sucrose content, 10 plants each from two tails were identified to make low- and high-sucrose bulk. In addition, data on yield contributing traits, *viz*., cane width (cm), cane length (cm), number of millable canes per row (NMCs), sucrose content (%), Brix value (%) and individual cane weight (kg) were also recorded. For all the phenotypic traits, analysis of variance (ANOVA) was based on the four check varieties as described by Gomez and Gomez (1984). The block effects for each of the agronomic traits were subtracted from the original mean individual data in order to get the adjusted mean for each trait for each genotype in both the years.

### Total RNA isolation and c-DNA library preparation for sequencing

The ten individuals to be used for bulking were selected from among the 187 F_1_ plants based on their over the years mean sucrose content and Brix value at 10 month growth stage, and accordingly tagged to make two (high and low sucrose) bulks. At the same time, RNA from the 8^th^ internode was isolated from the two parental lines and two bulks using Trizol reagent (Invitrogen). The quality of the total RNA was checked on denaturating agarose gel (1%), quantified using Qubit fluorometer (Thermo Fisher Scientific), and 100 µg RNA was sent to Xcelris Labs Limited (Ahmedabad, India) for Illumina sequencing (Illumina HiSeq 2000 Sequencing System, San Diego, USA).

Library preparation of the bulks was carried out after pooling equal amount of total RNA (100 µg) from each of the ten samples used for making the bulks. All the four samples (two parental lines and two bulks) were treated with NEBNext® Poly(A) mRNA Magnetic Isolation Kit (New England Biolabs, USA) to enrich mRNA from total RNA followed by fragmentation. The libraries were prepared using NEBNext® Ultra RNA Library Preparation Kit (New England Biolabs) as per manufacturer’s protocol. The fragmented mRNA was converted in to first-strand cDNA, followed by second-strand generation, poly(A)-tailing, adapter ligation, and finally ended by limited number of PCR amplification of the adaptor-ligated libraries. Library quantification and quality check was performed using DNA High Sensitivity Assay Kit. The amplified libraries were analyzed in Bioanalyzer 2100 (Agilent Technologies, Santa Clara, USA) using High Sensitivity (HS) DNA chip as per manufacturer’s instructions. The libraries were sequenced using 2×150 paired-end chemistry on Illumina HiSeq2000 Sequencing System (Illumina, San Diego, USA) for generating ∼10 GB of data per sample. The quality of the raw reads was checked using FastQC tool kit ver. 0.11.7 (Andrews, 2010). The raw reads were further trimmed using Trimmomatic Tool Kit (Bolger et al., 2014) and good quality paired-end reads were used for analysis.

### Sequence data analysis

The data analysis of sequence was carried out in two steps. First, in order to find out the differentially expressed genes (DEs) involved in sucrose accumulation, reads from all the samples were assembled *de novo* followed by mapping such reads back to the *de novo* assembly to get comparative read count. In the second step, the single nucleotide polymorphisms (SNPs) were identified directly from the structure of de Bruijn graphs constructed from the raw reads, analysed for differential enrichment and thereafter, the significantly differential SNPs were mapped on to the *de novo* assembly (Fig. 1).

**Fig. 1.**
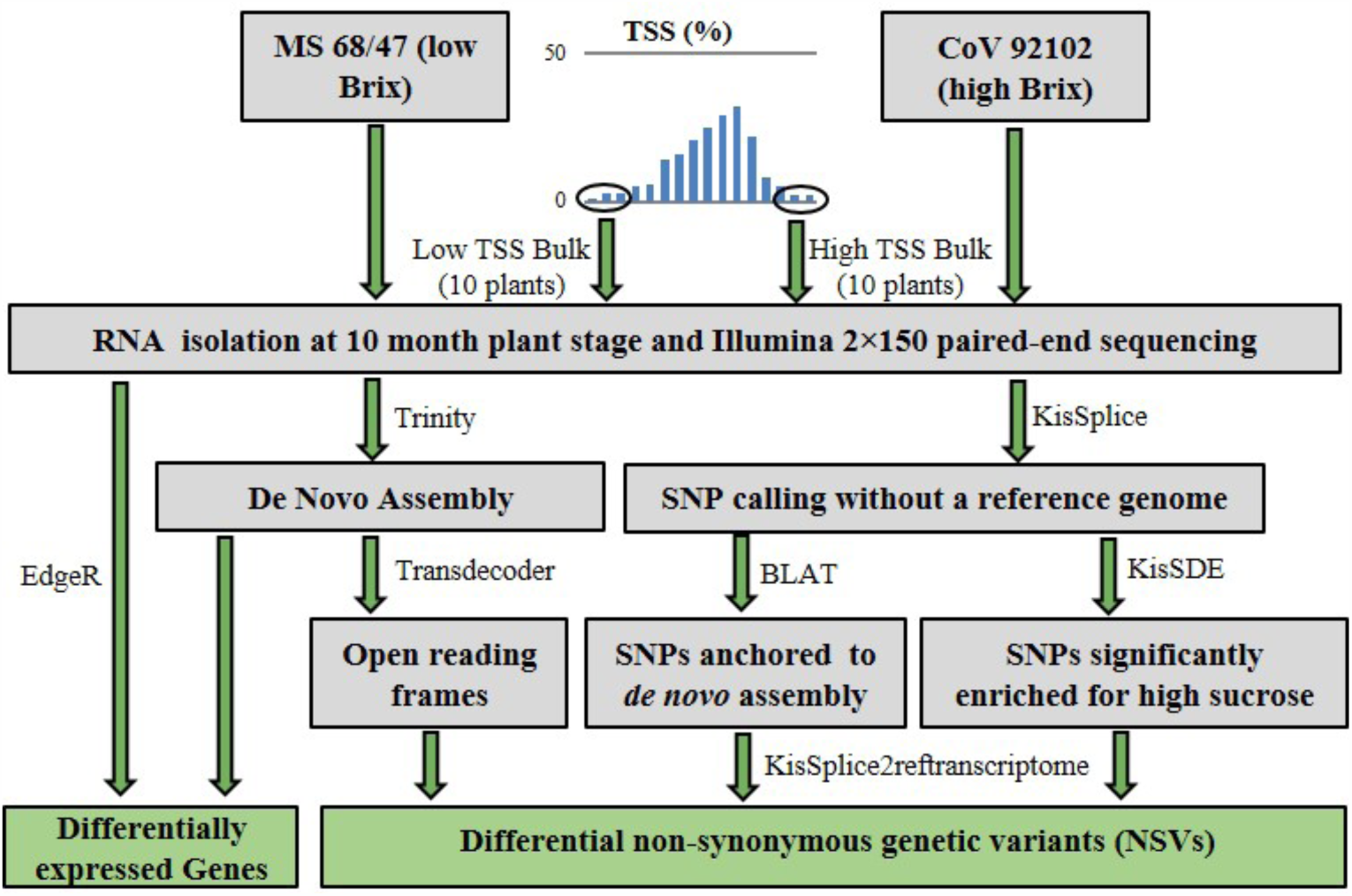
Schematic diagram of the materials and methods followed in this study. DEGs were identified by mapping the reads back to the *de novo* assembly. Differentially enriched NSVs were identified directly from the raw reads (using structures of the de Bruijn graph) and then anchored to the *de novo* assembly.

#### De novo assembly, and differential expression analysis

The raw reads were assembled *de novo* using De Brujin graph-based statistics (Trinity) on the web server TRUFA with default settings and by removing reads below 200 bp (https://trufa.ifca.es/web; Kornobis et al., 2015). In order to obtain InterPro annotations of the functional domains of the ‘genes’ as well as to assign gene ontology (GO) terms to the predicted genes, the *de novo* assembly was annotated using TRAPID (Rapid analysis of transcriptome data platform; http://bioinformatics.psb.ugent.be/webtools/trapid, Van Bel et al., 2013). Following this, Mercator web server was used to assign the sequences to functional bins (http://mapman.gabipd.org/app/mercator; Lohse et al. 2013).

In order to generate alignment files for obtaining read depths and identification of variants, the clean reads of all the four (two parents and high-low sucrose bulks) samples were mapped back to the *de novo* assembly using Bowtie 2 ver. 2.3.4.3 (Langmead and Salzberg, 2012). Read depths were measured using RSEM to get a measure of the expression level. Differential gene expression between the two parents was identified and re-confirmed in the bulks using the software EdgeR-Bioconductor Release 3.7 (Robinson et al., 2010). The common differentially expressed genes with more than 2-fold change and False Discovery Rate (FDR) corrected p-value of <0.005 were considered to be significantly differentially expressed (DE). These common DE genes were assigned to a functional bins using Mercator (Lohse et al. 2013) and visualized using MapMan (Thimm et al. 2004).

#### SNP variants calling and their mapping on de novo assembly

The criteria adopted for SNP calling in diploids are not applicable in polyploid crop like sugarcane due to confounding effects. Therefore, an alternate method of SNP calling was used in which SNPs were directly identified from the reads at the De Bruijn graph (DBG) construction phase (Lopez-Maestre et al. 2016), and thereafter anchored to the assembly using kisSplice package 2.4.0-p1-1 (Sacomoto et al. 2012). This approach is based on a theory that an SNP corresponds to an identifiable pattern, known as a bubble in the structure of a DBG that is built from two alleles (SNPs) of a locus (Peterlongo et al. 2010). Following this, the reads were mapped to each path of each bubble and frequency of the two variants was estimated. This facilitated reference-free identification of SNPs between the reads of the two parents as well as between the bulks. Thereafter, the SNPs were mapped on to the *de novo* assembly using BLAT (Kent 2002). The synonymous and non-synonymous SNPs were identified using Transdecoder ver. 5.3.0 (https://github.com/TransDecoder/TransDecoder). All the SNPs that were identified through kisSplice 2.4.0-p1-1 were tested for positive/negative association with early sugar accumulation capacity using kissDE package, which is based on testing a negative binomial distribution using a generalized linear model framework.

For quantifying the magnitude of effect of SNPs (Δf), the difference of allele frequencies (f) was calculated for each condition using the formula:

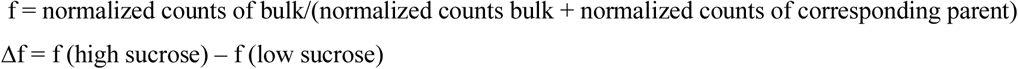

The Δf values for each SNP ranged between −1 (strong negative effect) and 1 (strong positive effect), while magnitudes close to 0 were identified as low effects on sucrose accumulation potential.

Following this, the contigs with differential non-synonymous SNPs (NSVs) present in the coding regions were assigned to a functional bins using Mercator and visualized using MapMan (Thimm et al. 2004) in order to visualize the quantitative effect of the SNV in the overall metabolic pathways.

## RESULTS

### Phenotyping

The parental lines (MS 68/47, CoV 92102) exhibited contrasting expressions for sucrose content, number of millable canes, cane diameter, and cane length and these traits segregated in their F_1_ population as well (Supplementary Table 2). There was a notable segregation within this F_1_ population for traits like sucrose content (17.44-8.9%), number of millable canes (1-17), cane length (110-320 cm), cane diameter (1.5-4.35 cm) and cane density (1.37-2.50 g/cc). Thereafter the ten genotypes possessing high sucrose (>16%) and ten having low sucrose (<10%) were selected for making high and low sucrose bulks.

### RNA-seq and *de novo* assembly

The RNAseq libraries of the two parental lines and two bulks generated 40-50 million paired-end reads in each sample. The mean size of library insert was 533 bp for MS 68/47, 554 bp for CoV 92102, while it was 511 bp for high sucrose bulk and 496 bp for low sucrose bulk. The high quality reads obtained after quality check, trimming and filtering of low quality unpaired reads were assembled *de novo* that resulted in a set of 183151 transcripts when all isoforms were considered. These transcripts represented 88939 assembled unigenes with a mean length of 783.11 bp, N50 value of 859 and GC content of 49.57%. The proportion of alignment of good quality paired-end reads for each of the four (CoV 92102, MS 68/47, high- and low-sucrose bulk) samples on the sequence assembly was >94%. The completeness of the *de novo* assembly measured on the basis of CEGMA alignment was 93.95% when only complete CEGs were considered and 100% when partial CEGs were also included (Table 1). Functional annotation using Mercator revealed that only 26.9% of the contigs could be assigned to functional bins, and of these, the largest (20%) proportion was in the ‘proteins’ bin followed by RNA (17%), signalling (11%) and transport (7%) (Supplementary Fig. 1).

**Table 1.** Raw reads and *de novo* assembly statistics of the sugarcane transcriptome using Trinity.

| Raw reads/sample | Total PE reads | Clean PE reads* | Reads aligned concordantly one time** (%) |
| --- | --- | --- | --- |
| MS/68/47 | 43529419 | 42497549 | 41.21 |
| CoV 92012 | 53013554 | 50866588 | 40.46 |
| High sucrose bulk | 52381343 | 50318499 | 45.45 |
| Low sucrose bulk | 46015104 | 44287834 | 39.42 |
| *De novo* assembly parameters | Genes | Transcripts |  |
| GC (%) | 49.57 |  |  |
| Number of genes/transcripts | 88939 | 183151 |  |
| N10 | 2914 | 2623 |  |
| N50 | 1215 | 859 |  |
| Average contig length | 783.11 | 588.41 |  |
| Complete CEGs count out of 248 (%) | - | 233 (93.95%) |  |
| Average orthologs/CEG | - | 2.57 |  |

\*Raw reads were cleaned and trimmed using Trimmomatic, \*\*Clean reads were aligned using Bowtie 2.

### Differentially expressed genes (DEGs) linked to sucrose accumulation

Differential gene expression was studied separately between the two parents, and between the two extreme bulks. The two parental genotypes had 15195 significant differentially expressed genes (DEGs) and the two bulks showed 7734 DEGs. A total of 2410 DEGs were identified that were common between the two groups (Supplementary Table 3, Fig. 2), and among them, 1559 were overexpressed during early sucrose accumulation, while 851 were overexpressed in late sucrose accumulation conditions. A majority of these DEGs were involved in protein activation, synthesis and degradation, while, a significant number was involved in signalling, stress and transport (Supplementary Table 3). Under high sucrose accumulating conditions, a very high degree (>10-fold) of upregulation was noted in the genes involved in sugar and metabolite transport at mitochondrial membrane, cell vesicle transport, etc. (Table 2). A lot of structural variation within the genes was noted, in addition, there was up- as well as down-regulation of alternative forms of the same gene. For example, structural variants of Cytochrome P450 and UDP-glycosyl transferase exhibited contrasting differential expression between high- and low-sucrose accumulation conditions (Supplementary Table 3).

**Table 2.**
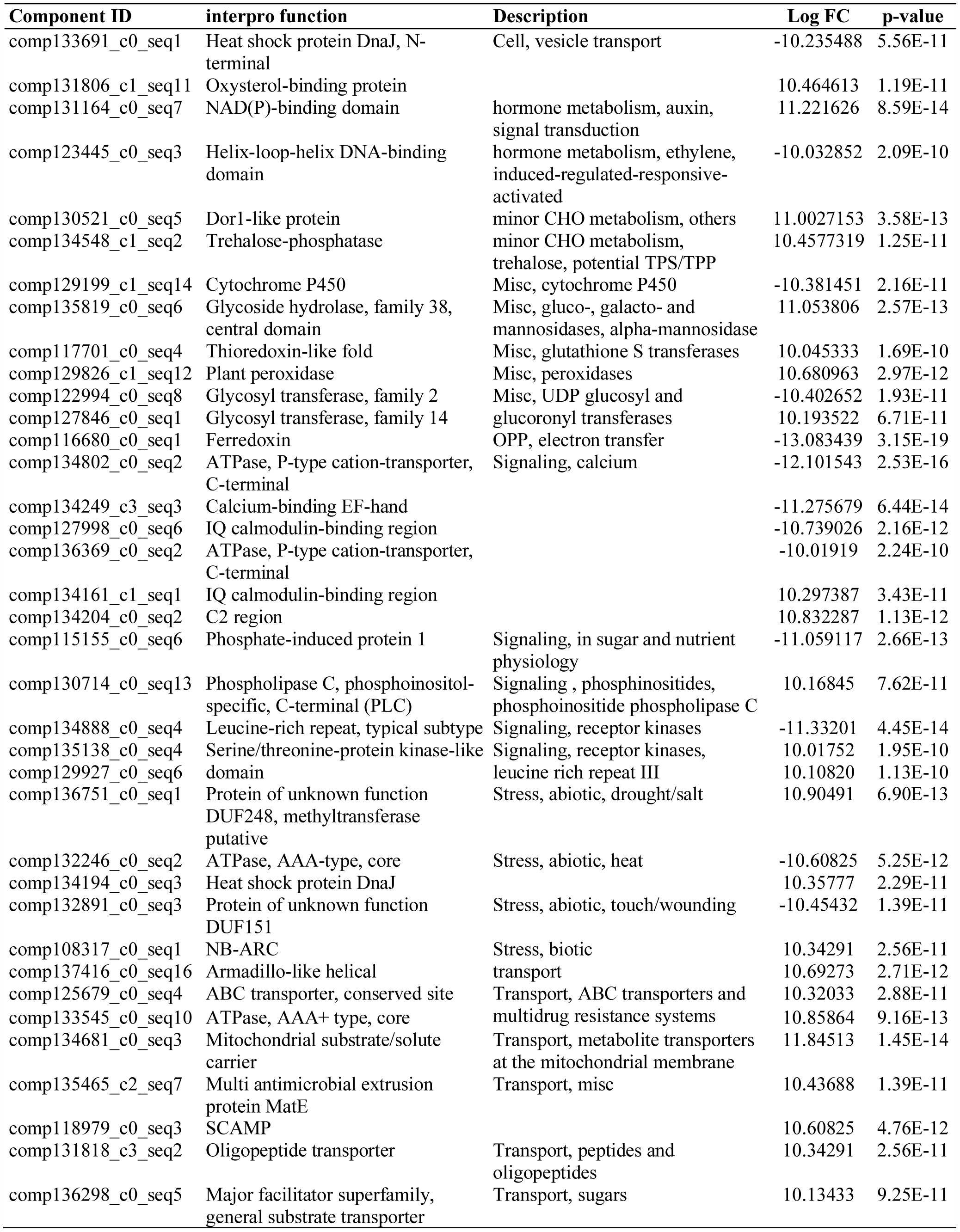
Differentially expressed genes during high sucrose accumulation conditions identified in this study. Only those genes are listed that showed >10-fold up- or down-regulation).

#### Overall metabolic pathway

When the overall metabolic pathway was visualized on MapMan, it was found that that genes for cellulose and hemicellulose synthesis were upregulated during early sucrose accumulation (Fig. 3). Similarly, genes related to minor carbohydrate metabolism (trehalose, inositol phosphatase, potential TPS/TPP) were also upregulated during early sucrose accumulation. In addition, a number of genes related to cell wall degradation, polygalacturonases, phospholipid synthesis, lipid degradation, starch and sucrose synthesis (AGPase and FBPase) were upregulated. Whereas, genes associated with starch degradation, *e*.*g*., glucan water dikinase and ISA3 were downregulated. Another significant gene producing no apical meristem (NAM) protein exhibited ten-fold upregulation during early sucrose accumulation.

**Fig. 2.**
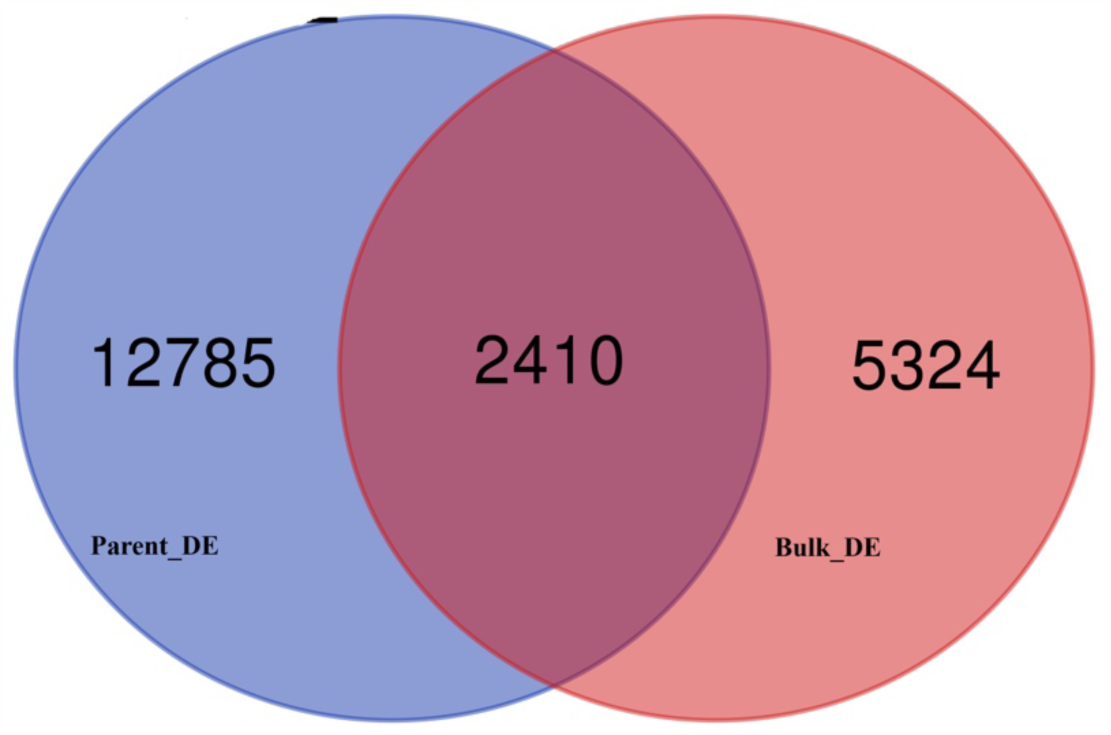
Venn diagram of differentially expressed genes (DEGs) between the two parents (MS 68/47, CoV 92102) and between the high and low sucrose bulks.

**Fig. 3.**
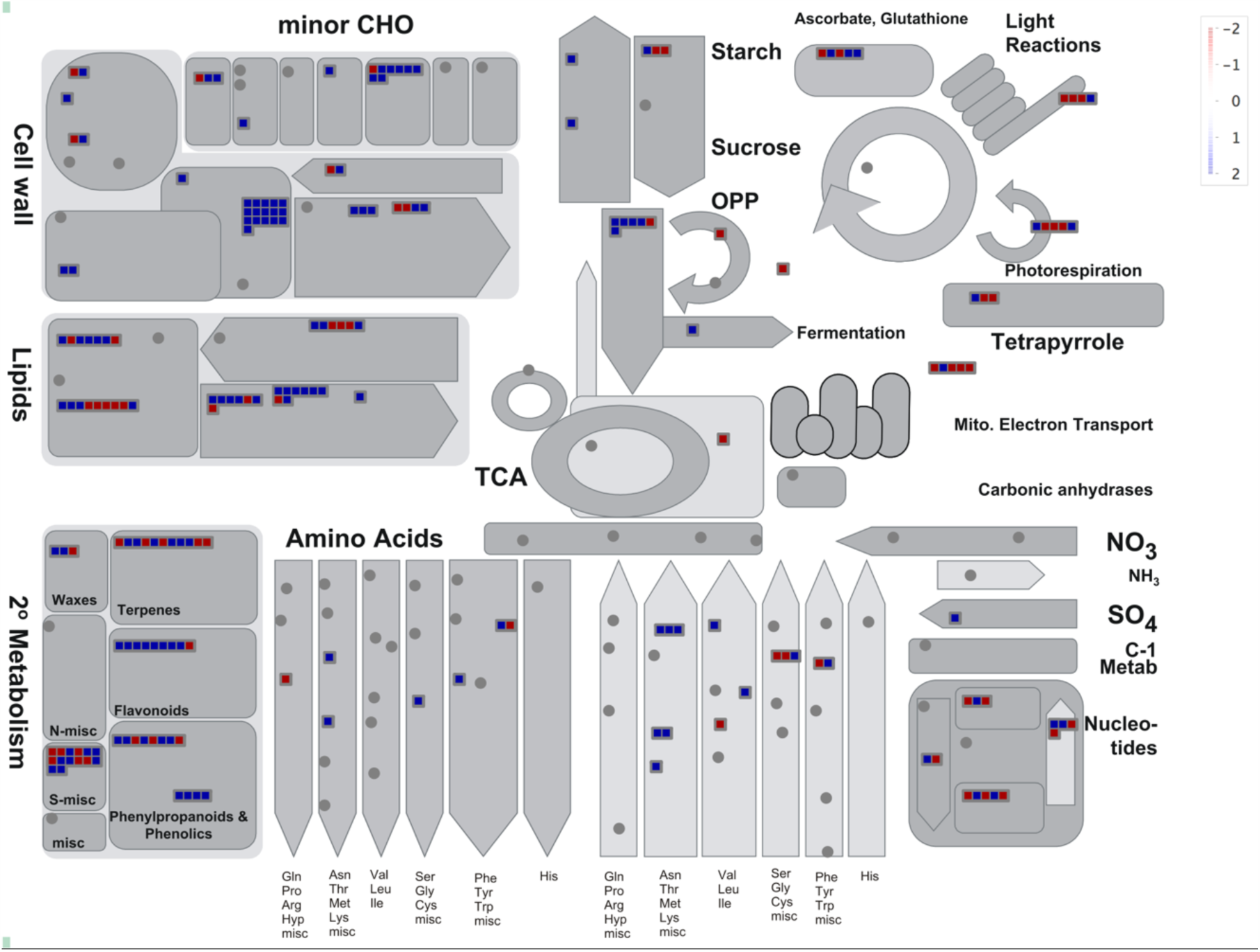
A metabolic pathway overview of the common DEGs between the two parents and two extreme bulks. The expression level of each gene in the pathway has been represented as blue for the highest expression and red for the lowest expression.

#### Sucrose metabolism

Sucrose phosphate synthase (SPS; EC 2.4.1.14), sucrose phosphate phosphatase (SPP; EC 3.1.3.24), invertase (EC 3.2.1.26), sucrose synthase (SuSy; EC 2.4.1.13), fructokinase (EC 2.7.1.4) and hexokinase (EC 2.7.1.1) are the key enzymes involved in sucrose metabolism. When the variation was considered only between high- and low-sucrose parents, it was observed that SPS III was upregulated eight-folds and SPS V was downregulated two and a half-folds (Fig. 4a), and rest of the sucrose pathway enzymes (SuSy, invertase, fructokinase and hexokinase) which are mainly involved in the further breakdown of sucrose were predominantly downregulated. In addition, the alternate isoforms of neutral invertase, fructokinase and hexokinase were identified that were upregulated in the high sucrose parent. Similarly, when the two bulks were compared, a 10-fold downregulation of SPS III during early sucrose accumulation was noted (Fig. 4b). In the case of SuSy as well as hexokinase, of the two isoforms they had, one was upregulated while the other one was downregulated. Both neutral and vacuolar invertases were upregulated in the high sucrose bulk (Fig. 4). But an overlap of differential regulation could not be observed for various enzymes involved in sucrose metabolism when parents and the bulks were considered together.

**Fig. 4.**
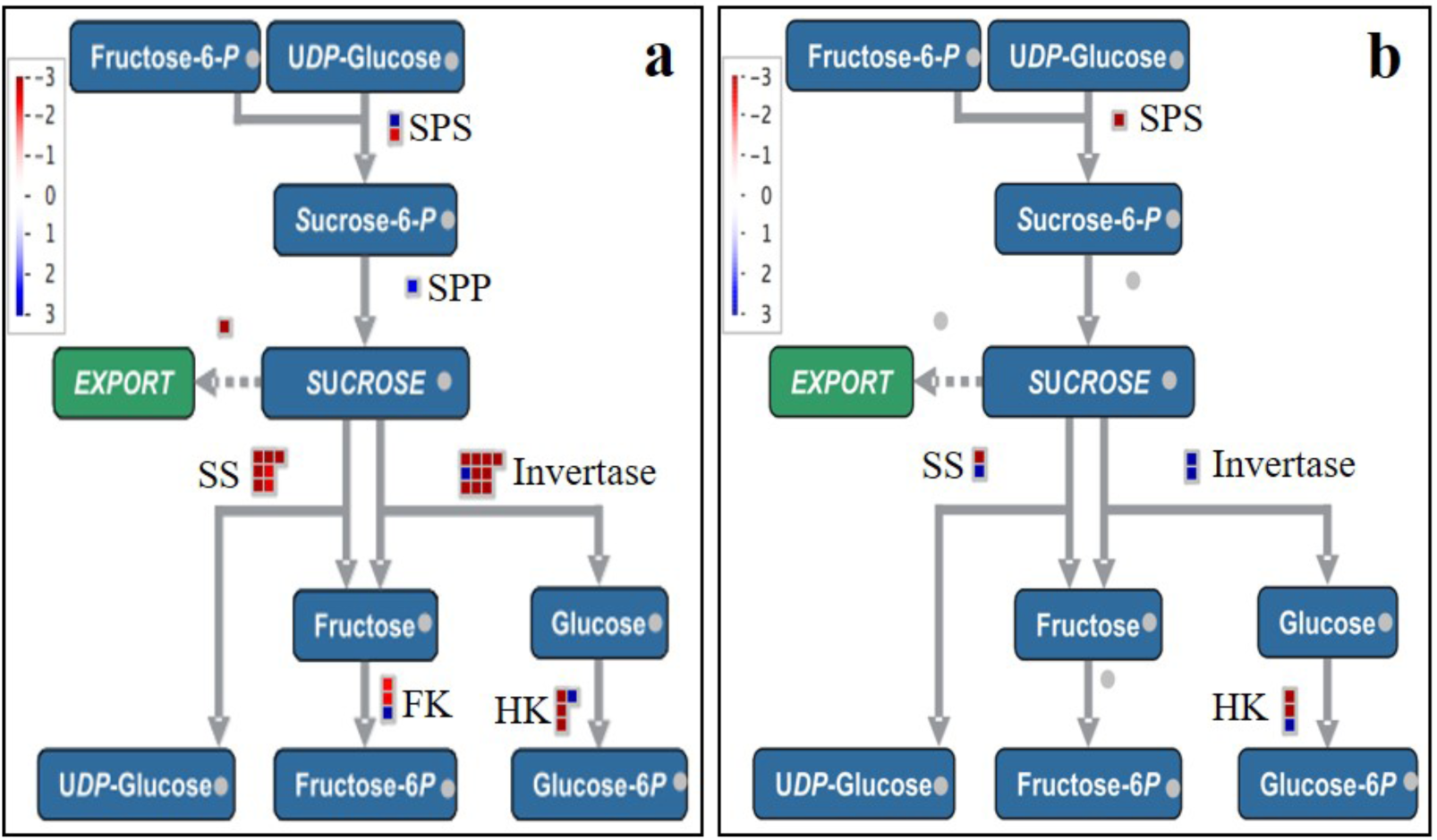
Differential gene regulation of sucrose synthesis pathway by differentially expressed genes between the high and low sucrose parents (a), and between the high and low sucrose F_1_ bulks (b).

#### Glycolysis and photosynthesis

Barring pyruvate kinase which was downregulated, all other genes involved in glycolysis, *viz*., fructose 2,6 bisphosphatase, phosphoglycerate mutase, phosphoenolpyruvate kinase and phosphoenolpyruvate carboxylase kinase were upregulated (Fig. 3, Supplementary Table 3). Most of the genes associated to photorespiration, mitochondrial electron transport and oxidative photophosphorylation were downregulated during early sucrose accumulation, while the PS II-associated genes were downregulated (Fig. 3, Supplementary Table 3).

#### Transcription factors

Among the transcription factor (TF) family genes, Histone DAse, Histone, WRKY, JUMONJI, bZIP, Homeobox TFs, MADS box, MYB-related TFs, C3H TFs, zinc-finger family, ELF3 and a silencing group TF was upregulated during high sucrose accumulation. Whereas, TFs, *viz*., Histone acetyltransferase, ARR-B, Constans-like zinc finger family (C_2_C_2_-CO like), C_2_H_2_ zinc finger family and an orphan TF involved in DNA repair were down regulated (Supplementary Table 3).

### Non-synonymous genetic variations (NSVs) linked to high sucrose accumulation

A total of 27626 SNPs with a very good coverage (>100) in the parents as well as the bulks, that had been extracted directly from the De Bruijn graphs and satisfied the FDR criterion (P <0.005) were selected (Supplementary Table 4). Of the 27626 SNPs identified, the maximum SNPs were located in genes coding for protein synthesis and degradation, followed by signalling and transport. Notably, there were some genes that showed differential regulation only at the SNP level and not in the form of DEGs, and we were able to spot such differential regulation visible only at the allelic level in case of gluconeogenesis, fermentation and C1 metabolism. Moreover, the number of variable SNPs detected was much higher than the number of DEGs and a higher degree of allelic variation was observed in minor and major carbohydrate metabolism, Glycolysis, TCA, mitochondrial and electron transport genes (Table 3). All the metabolic pathways exhibited alternative forms of genes/alleles that were up- and down-regulated. Of the identified SNPs, 21511 were present in the coding region of the genes, and among them 9905 were non-synonymous variants (NSVs). Due to the fact that this model could quantify the individual effect of each SNP on trait expression, following the 9905 NSVs located in the coding regions of the genes, and visualizing them in different cellular pathways, it became possible to identify important allelic variations of genes that were differentially enriched during early sucrose accumulation (Supplementary Table 4). A total of 43 NSVs had a Δf value >0.5, and such NSVs are primarily located on the genes involved in hormone metabolism, mitochondrial electron transport, signalling, transport and TCA, etc. (Table 4).

**Table 3.** Frequency distribution of differentially expressed genes and NSVs into functional bins as defined by Mercator.

| Functional bin | DEG | Differential NSV |
| --- | --- | --- |
| Amino acid metabolism | 20 | 688 |
| Biodegradation of Xenobiotics | 1 | 75 |
| C1-metabolism | - | 67 |
| Cell cycle, division and organization | 55 | 919 |
| Cell vesicle transport | 12 | 366 |
| Cell wall | 31 | 510 |
| Co-factor and vitamin metabolism | 5 | 98 |
| Cytochrome P450 | 16 | - |
| Development | 51 | 838 |
| DNA synthesis and repair | 75 | 538 |
| Fermentation | 1 | 102 |
| Gluconeogenesis | - | 27 |
| Glycolysis | 5 | 233 |
| Hormone metabolism | 40 | 550 |
| Lipid metabolism | 38 | 634 |
| Major carbohydrate metabolism | 5 | 155 |
| Metal handling and storage | 5 | 133 |
| Micro RNA | - | 6 |
| Minor carbohydrate metabolism | 14 | 138 |
| Mitochondria/electron transport | 5 | 202 |
| N-metabolism | 1 | 24 |
| Nucleotide metabolism | 13 | 205 |
| OPP | 2 | 89 |
| Oxidases | 11 | - |
| Peroxidases | 5 | - |
| Photosynthesis | 6 | - |
| Polyamine metabolism | 1 | 37 |
| Protein activation, synthesis and degradation | 200 | 4681 |
| Protein targeting | 12 | 571 |
| Redox | 8 | 334 |
| RNA processing | 38 | 543 |
| RNA regulation of transcription | 151 | 2732 |
| Secondary metabolism | 31 | 423 |
| Signaling | 77 | 1287 |
| Stress | 68 | 757 |
| TCA | 1 | 217 |
| Tetrapyrrole synthesis | 3 | 42 |
| Transport | 70 | 1212 |
| Transport, sugars | 10 | - |
| UDP-glycosyl transferase | 20 | - |
| Not assigned | 1256 | 94194 |

**Table 4.**
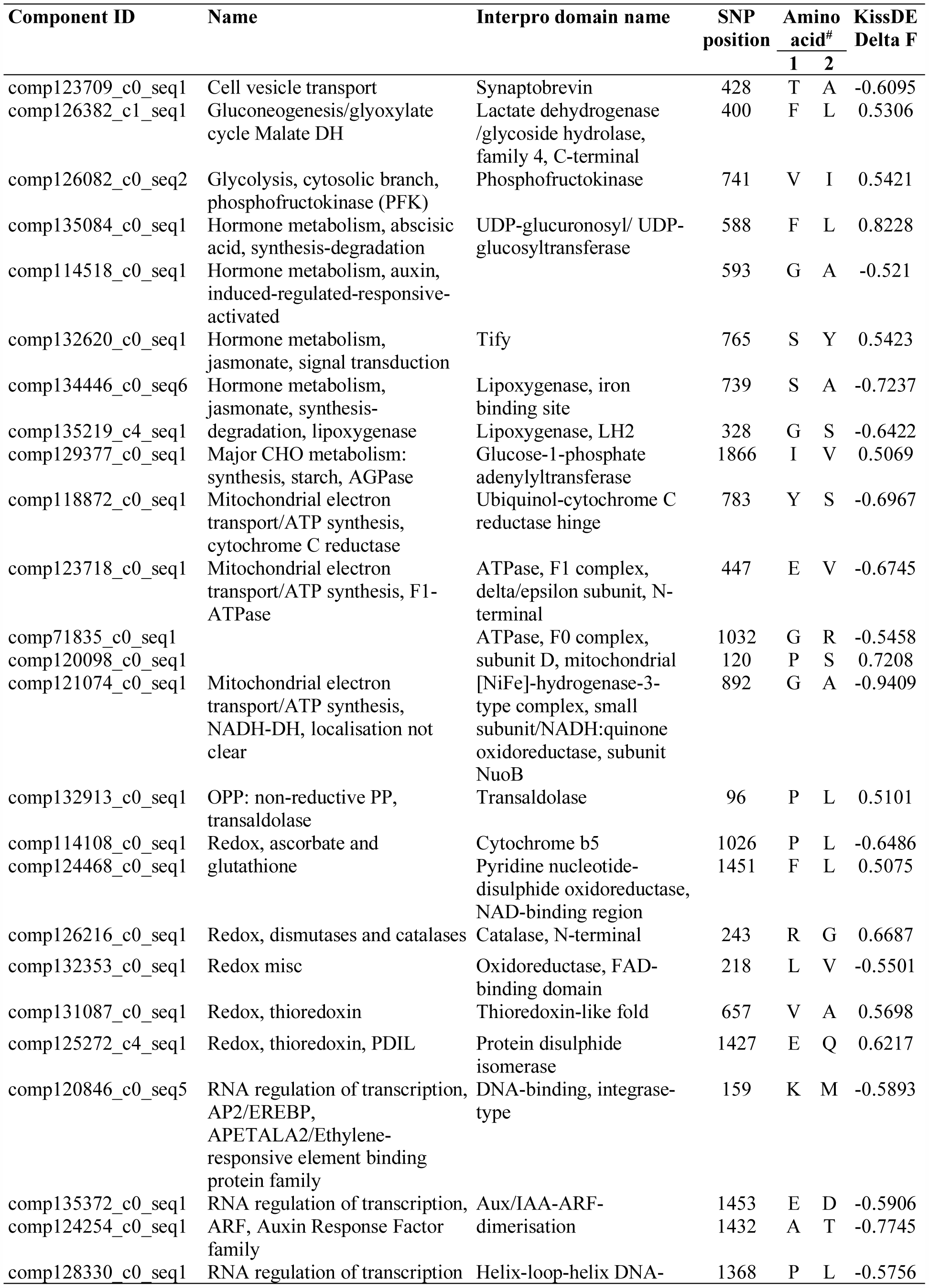
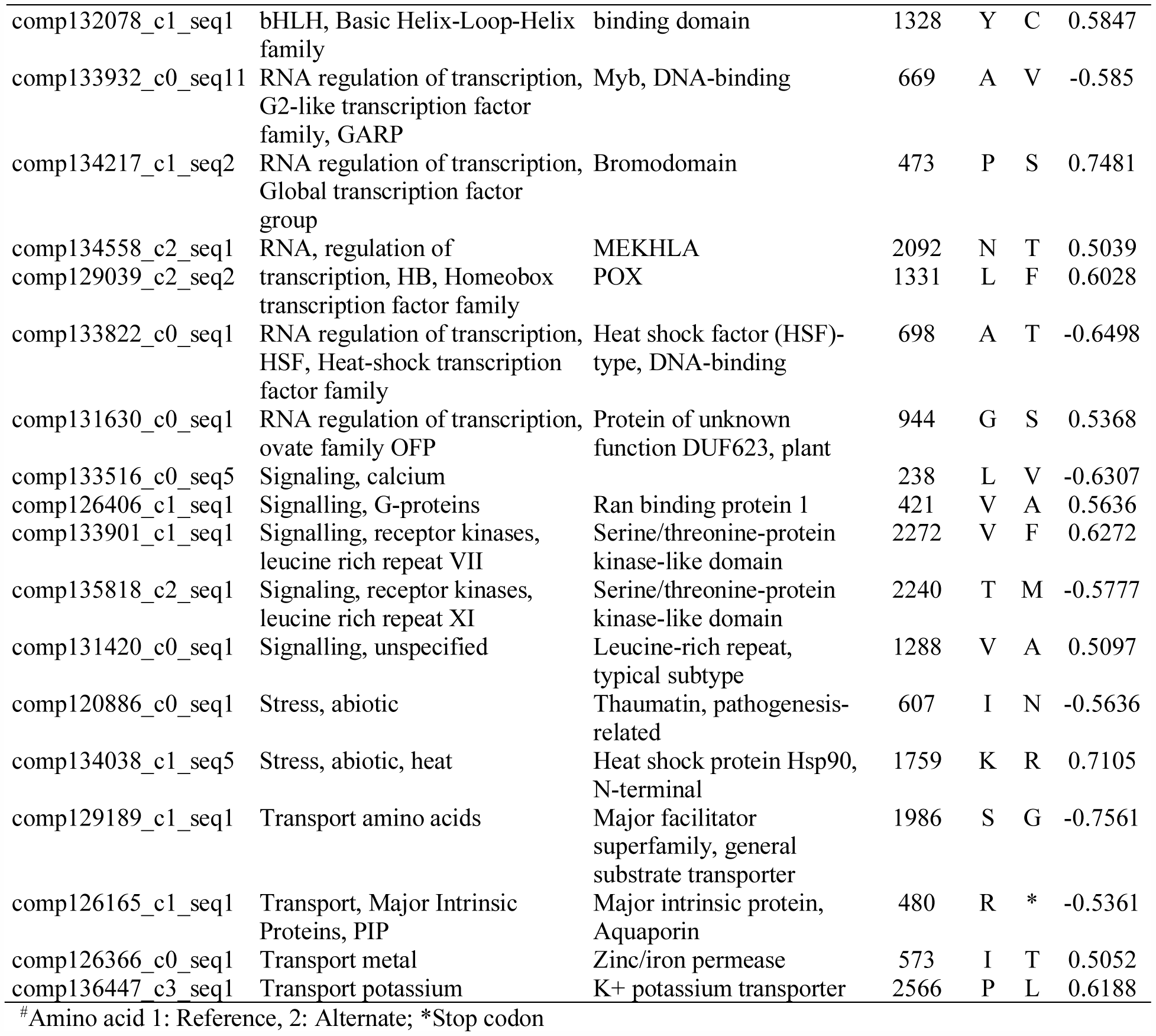
Highly significant (Δf > 0.50) differentially enriched non-synonymous variants linked to sucrose accumulation.

#### Overall metabolic pathway

The upregulation of genes associated with minor carbohydrate metabolism pathways was more explicitly visible at the allelic level since most of the genes (Raffinose synthase, Trehalose synthase, Trehalose phosphate, FGGY-type carbohydrate kinase, myoinositol oxygenase, myoinositol phosphatase, xylulose kinase and glycosyl transferase) of this pathway exhibited differential NSVs during early sucrose accumulation. In contrast, majority of the differential NSVs in the genes coding for cell wall proteins, like hydroxyproline rich glycoproteins (HRGPs) showed a negative effect NSVs for early sucrose accumulation. Among the genes coding for major enzymes of carbohydrate metabolism, the differential NSVs favouring early sucrose accumulation were detected in starch synthase, β-amylase and *AGPase* genes. In contrast, genes coding for α-amylase, starch branching enzyme, starch phosphorylase, starch D enzyme had mostly differential NSVs with negative effect on early sucrose accumulation. In the case of glutathione ascorbate cycle, that plays a role in the H_2_O_2_ detoxification, majority of genes had SNPs that showed a negative effect on early sucrose accumulation. Glucose-6-phosphate dehydrogenase and transketolase of oxidative pentose phosphate pathway had a positive effect differential NSVs, whereas, negative effect NSVs were detected for 6-phosphogluconolactonase, 6-phosphogluconate dehydrogenase and ribulose-phosphate-3-epimerase. Some genes like transaldolase were detected in two of their isoforms, one that had a positive effect differential NSV and other had a negative effect differential SNP. Majority of the fermentation-associated genes (*LDH, PDC, ADH* and aldehyde dehydrogenase) possessed a positive effect NSV. Almost all the sulphur assimilation genes, *viz*., *ATPS, ARP*, sulphite redox and sulphite oxidase had positive effect NSVs. All the genes associated with wax metabolism had a negative effect NSV for sucrose accumulation (Fig. 5).

**Fig. 5.**
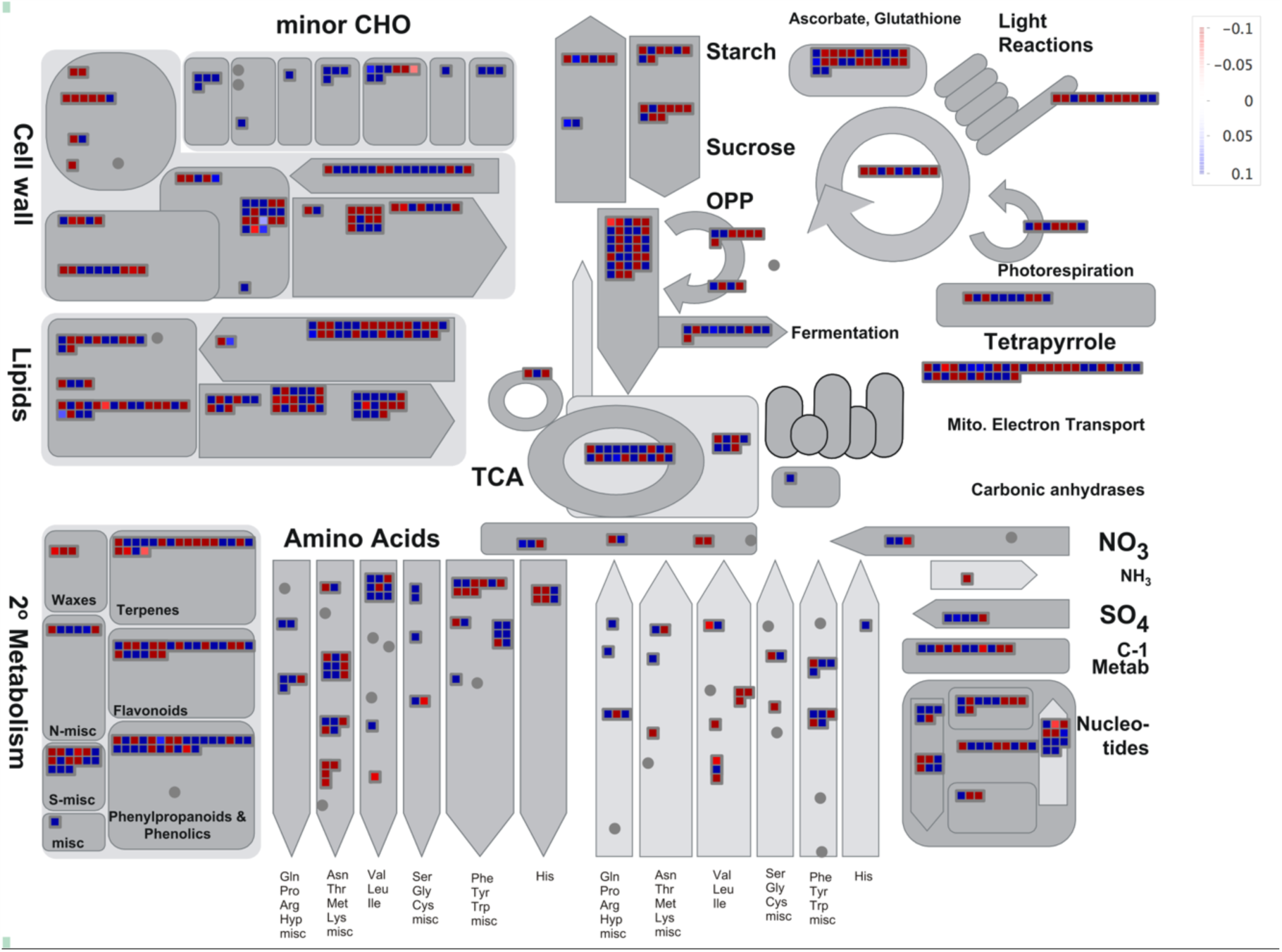
A metabolic pathway overview of the differential NSVs between the high and low sucrose accumulation conditions. The expression level of each NSV in the pathway (measured as Δf value) has been represented as blue for the highest enrichment and red for the lowest enrichment.

#### Sucrose metabolism

Interestingly, among the major sucrose metabolism enzymes (SPS, SPP, SuSy, hexokinase, fructokinase and invertase), differential NSVs were identified that had both negative as well as positive effect on early sucrose accumulation. In the case of SPS, five NSVs were detected within the glycosyl transferase domain, but no NSV was detected in the SPP-binding S6PP domain. The SPP on the other hand, showed presence of just one NSV in its S-6PP domain. The two isomers (SuSy I, SuSy II) of sucrose synthase exhibited few differential NSVs; SuSy I had one differential NSVs in its RfaB domain. Sucrose synthase II showed three NSVs in the S6PP domain located within its active site. Similarly, a number of very important functional NSVs were detected within the coding regions of other enzymes, *viz*., neutral/alkaline invertase, acid invertase, soluble starch synthase starch branching enzyme, hexokinase and 1,4 α-glucan branching enzyme of the sucrose-starch metabolic pathway (Fig. 6; Table 5).

**Table 5.** Differentially enriched non-synonymous variants identified on the genes involved in sucrose metabolism.

| Component ID | Description | SNP position | Delta F value | p-value | Amino acid<sup>#</sup> |  |
| --- | --- | --- | --- | --- | --- | --- |
|  |  |  |  |  | 1 | 2 |
| comp132954_c0_seq1 | Chloroplastic/amyloplastic 1,4-alpha-glucan-branching enzyme | 717 | -0.294 | 0.003412 | R | H |
|  |  | 1015 | 0.1385 | 0.000713 | D | N |
|  |  | 2701 | 0.1476 | 0.001178 | R | W |
|  |  | 576 | 0.1652 | 0.000329 | G | V |
|  |  | 1127 | 0.1881 | 0.003732 | R | S |
|  |  | 780 | 0.4219 | 5.07E-10 | S | F |
| comp135650_c0_seq1 | Chloroplastic/amyloplastic soluble starch synthase 1 | 1269 | 0.3232 | 8.16E-05 | A | G |
| comp124275_c1_seq1 | Hexokinase 1 | 314 | -0.1159 | 1.01E-05 | Q | E |
|  |  | 330 | 0.3544 | 9.58E-05 | V | A |
|  |  | 624 | -0.1232 | 0.000233 | V | M |
| comp123096_c0_seq1 | Neutral/alkaline invertase | 1314 | -0.1361 | 0.000292 | S | N |
|  |  | 1991 | 0.1945 | 0.002676 | I | V |
| comp136607_c3_seq1 | Soluble acid invertase 9 | 642 | -0.2582 | 4.57E-05 | G | D |
| comp136607_c0_seq1 |  | 483 | -0.1641 | 1.72E-06 | G | A |
|  |  | 868 | 0.1469 | 0.001066 | E | D |
| comp133566_c0_seq1 | Starch branching enzyme | 2397 | -0.2405 | 5.43E-05 | S | P |
|  |  | 2573 | 0.4272 | 0.000368 | V | A |
| comp126489_c0_seq1 | Sucrose phosphate phosphatase | 1300 | 0.3026 | 7.53E-07 | T | S |
| comp134759_c1_seq1 | Sucrose phosphate synthase | 2220 | -0.0887 | 0.000234 | F | L |
|  |  | 1419 | 0.0825 | 2.14E-07 | R | C |
|  |  | 452 | 0.1463 | 2.36E-05 | S | N |
| comp131008_c0_seq1 | Sucrose synthase 1 | 692 | -0.1201 | 0.002494 | Q | R |
| comp124821_c0_seq1 | Sucrose synthase 2 | 2512 | -0.2848 | 0 | N | S |
|  |  | 639 | 0.1319 | 0 | E | D |
|  |  | 1292 | 0.1396 | 0 | H | D |

<sup>#</sup>Amino acid 1: Reference, 2: Alternate

**Fig. 6.**
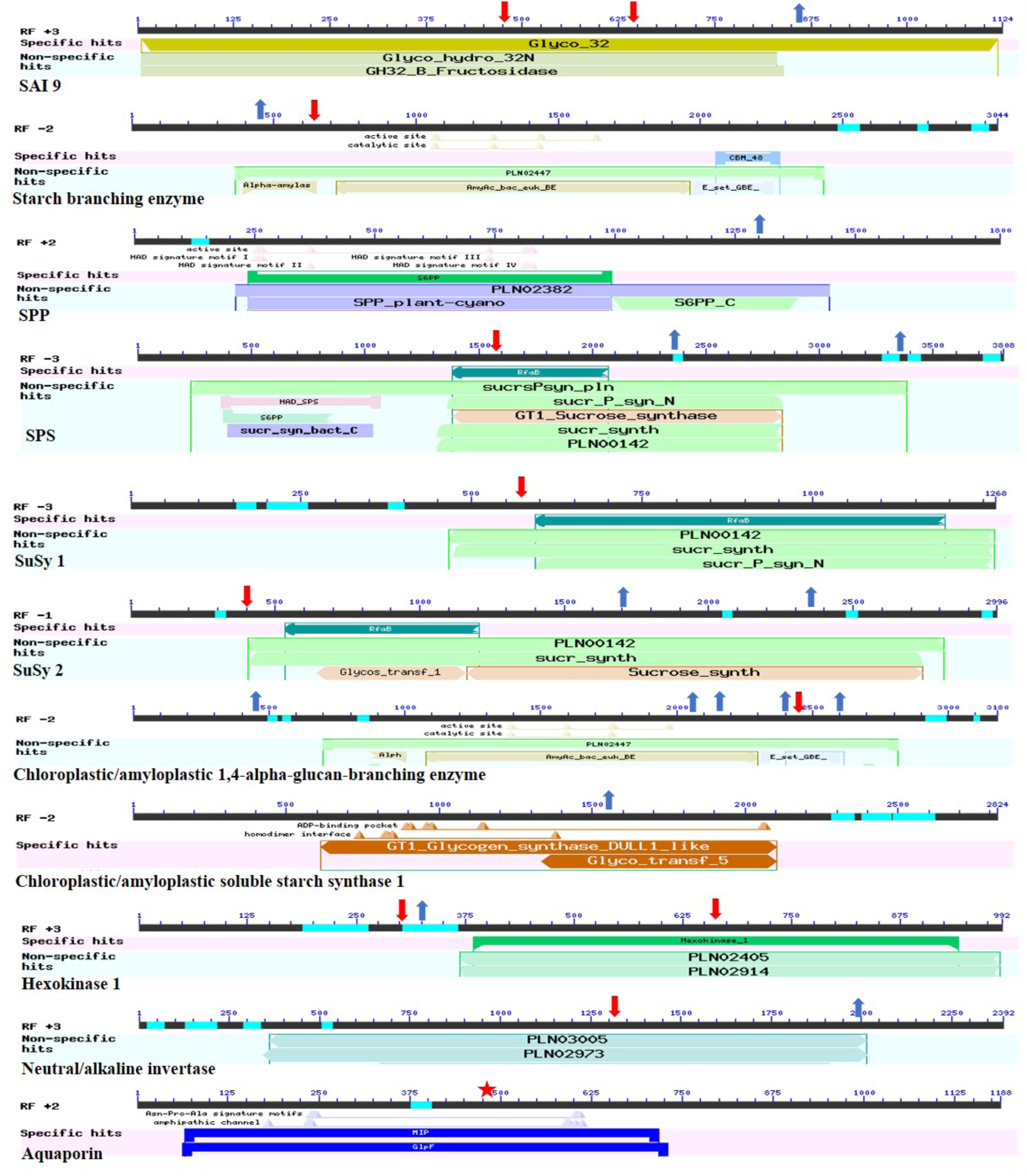
Location of differentially enriched NSVs on the genes involved in the sucrose metabolism pathway. A blue arrow indicates a positive effect NSV, while a red arrow indicates a negative effect on the trait. (SAI: Soluble acid invertase, SPP: Sucrose phosphate phosphatase, SPS: Sucrose phosphate synthase, SuSy: Sucrose synthase).

**Fig. 7.**
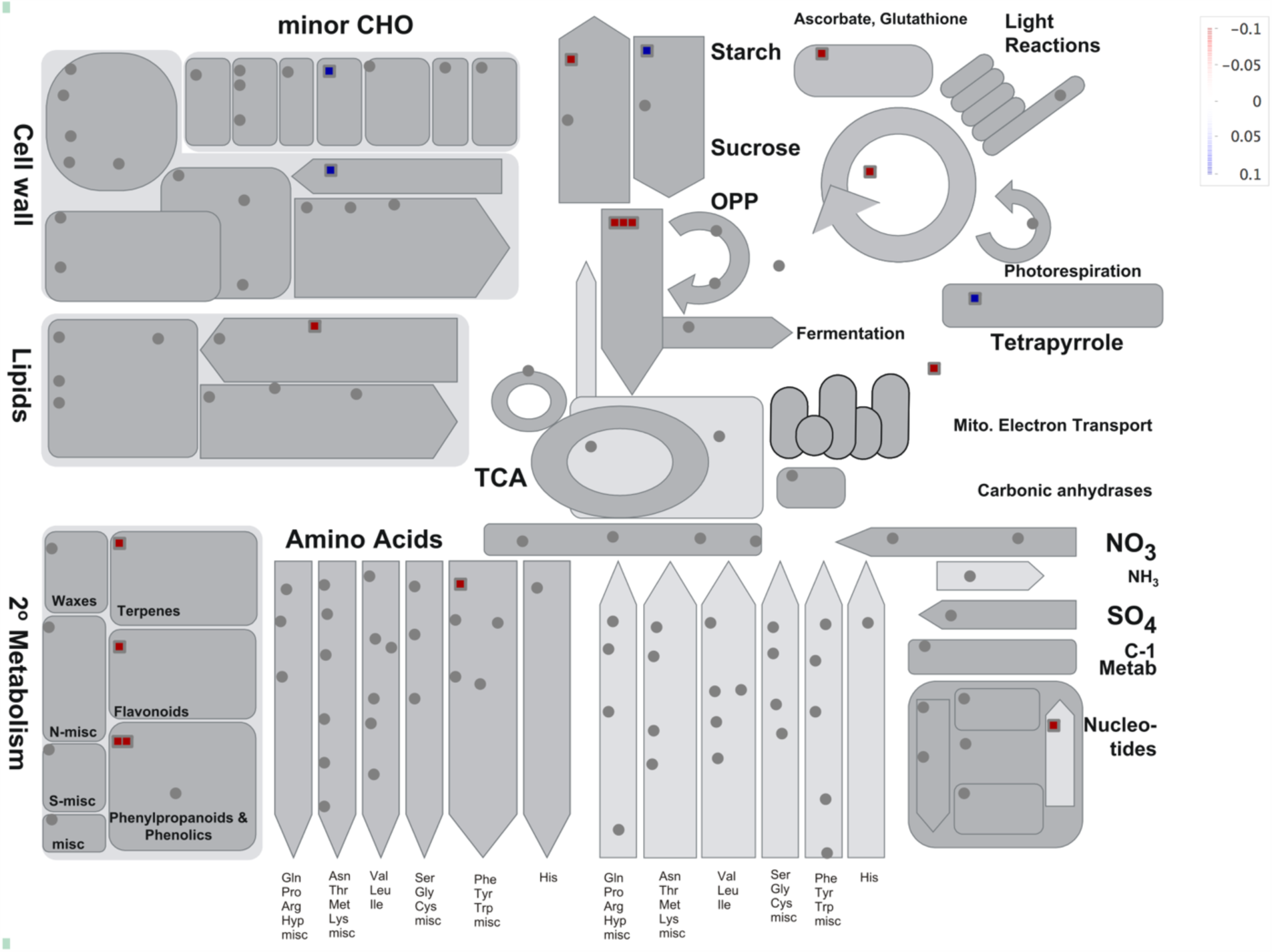
A metabolic pathway overview of the common premature stop codons between the two parents and two extreme bulks. The expression level of each gene in the pathway has been represented as blue for the highest expression and red for the lowest expression.

#### Photosynthesis and mitochondrial electron transport chain

In the case of light reaction of photosynthesis, there were favourable as well as unfavourable differential SNPs in the genes coding for photosystem I (PS I) and II (PS II) polypeptide subunits. For example, in the case of PS I, LHC-1 differential SNPs had negative effects; such SNPs were also found in genes coding for cytochrome b6/f and ATP synthase. In the case of state transition genes, two positive and one negative effect SNPs were detected. Among the genes involved in Calvin cycle, NSVs with negative as well as positive effect were detected in aldolase and Rubisco. Positive effect SNP was detected in transketolase, while negative effect SNP was detected in genes coding for TPI and sedoheptulose bisphosphatase (Supplementary Table 4, Fig. 5).

In the mitochondrial electron transport chain, there were positive and negative effect SNPs in NADH-dehydrogenase complex I, carbonic anhydrase, cytochrome C reductase, cytochrome C, Cytochrome C oxidase and F1-ATPase. Electron transfer flavoprotein had only a negative effect SNP. The nitrate metabolism genes glutamate synthase and glutamine synthetase had a predominant positive effect SNP (Supplementary Table 4, Fig. 5).

#### Glycolysis and TCA

In context to glycolysis enzymes, both positive as well as negative effect differential NSVs were identified for aldolase, phosphoglycerate mutase, phosphoenolpyruvate carboxylase, phosphofructokinase, pyruvate kinase and pyrophosphate-fructose-6-P-phosphotransferase. The positive effect NSVs were present in genes coding for triose-phosphate isomerase (TPI), enolase and phosphoenol pyruvate carboxylase kinase, whereas, SNPs that had a negative effect on the trait were detected in genes coding for phosphoglucomutase, glucose-6-phosphate isomerase, glyceraldehyde-3 phosphate dehydrogenase (GAPDH) and phosphoglycerate mutase. Similarly, in the case of the TCA cycle genes, there were positive as well as negative effect NSVs in pyruvate dehydrogenase, aconitase, IDH, 2-oxalogluterate dehydrogenase, succinyl Co-A ligase and succinate dehydrogenase. The only positive effect NSVs were detected for carbonic anhydrase and malate dehydrogenase (Supplementary Table 4, Fig. 5).

#### Transcription factors

Among the transcription regulation factors, the positive effect NSVs were located in TFs of families such as ARR, GARP, Triple-Helix, Global transcription factor group, Histone core H3, Heat shock, C2C2(Zn) Dof zinc finger, GATA, Auxin/IAA, methyl binding domain proteins, NIN-like bZIP-related, MYB-related, Pseudo ARR and chromatin remodelling factors. The TFs with predominant negative effect NSVs were from basic-helix-loop-helix, Alfin-like, HDA, EINB-like (EIL), H2A, H2B, histone, PWWP domain protein, C3H zinc finger and DNA methyltransferases (Supplementary Table 4).

#### NSVs resulting in stop codons

In addition to the positive/negative effect NSVs detected in different genes, a set of 28 NSVs resulting in non-sense mutations (premature stop codons), which could possibly result in truncated proteins upon translation were also detected. These non-sense mutations were visualized on MapMan (Supplementary Table 4, Fig. 7), and the allele frequency of the NSVs resulting in stop codons ranged from 4 to 100%. A very important negative effect premature stop codon was identified within the amphipathic channel domain of gene coding for a water channel protein, aquaporin (Fig. 6; Table 4). Of the 28 such NSVs, nine were tagged to transcripts that could be annotated by InterPro. The presence of premature stop codons in the coding regions of genes for PIP, aldolase, MYB-related transcription factor family, ubiquitin, CRR1 showed a negative effect on the trait. On the other hand, the presence of a stop codon in genes coding for auxin hormone metabolism, SET domain transcription regulation family, serine protease, thioredoxin, etc. had a positive effect on the trait. However, most of these stop codons were linked to transcripts that could not be annotated.

## DISCUSSION

Sugarcane is a long duration crop and there exists a significant variation in agronomical traits including the maturity time and sucrose content among different varieties. Identification of the source of this variation would be helpful to create short-duration varieties with high sucrose accumulation potential. In highly polyploid crops, allelic variation and enrichment plays a major role in the trait expression. Considering advantages like low sequencing cost, identification and quantification of allelic variation and enrichment, RNAseq in recent past has become a better alternative to DNA sequencing based studies. In sugarcane, there have been a few studies using transcriptome sequencing of bulks contrasting for sucrose (Thirugnanasambandam et al., 2017, 2019) and fibre content (Kasirajan et al., 2018), which led to identification of important differentially expressed genes associated with such traits. Most of the previous studies on RNA-seq in sugarcane have focussed on sucrose accumulation at mid-late stage, but to our knowledge, this is the first report in which triggers of early sucrose accumulation were also targeted.

SNP calling in polyploids is confounded, and hence the usual criteria adopted for diploids are not applicable. In diploids, SNP calling can easily be done by mapping the reads to the genome assembly for variant calling, but due to the polyploidy and complexities of sugarcane genome, an alternate method was adopted, wherein, SNPs were directly identified from the reads at the De Bruijn graph construction phase, and thereafter anchored to the assembly (Lopez-Maestre et al. 2016). This method is based on a concept of Peterlongo et al. (2010) and Iqbal et al. (2012) that a SNP corresponds to an identifiable pattern, known as a ‘bubble’ in the structure of a De Bruijn graph. Hence a De Bruijn graph that is built from two alleles (SNPs) of a locus will correspond to a ‘bubble’ on the graph. Following this the reads were mapped to each path of each ‘bubble’ and frequency of the two variants was estimated. Adoption of such an alternate method in present study led to significant differential allele enrichment under the two contrasting conditions. Brum (2018) suggested that in sugarcane, the use of allelic signal ratio could be better for predicting genotype since it eliminates the essentiality of complete reference genome sequence and exact ploidy knowledge. In addition, since alleles are screened on the basis of their abundance in different conditions, their relative effect on the trait could be also be measured. The SNPs within the coding region may cause a change of a single amino acid (non-synonymous variations; NSV), or may not effect a change in amino acid due to degeneracy of the genetic code (synonymous). As a result, these NSVs could have a significant positive/negative impact on the overall structure and function of protein. NSVs causing premature stop codons could lead to protein truncation which could lead to transcript degradation and negative influences of the protein (Nagy and Maquat, 1998). In the present study, the 28 pre-mature stop codons were located in genes involved in significant steps of the metabolic pathways and exhibited both positive as well as negative effect on the trait. Fernald et al. (2011) and Shendure and Akey (2015) speculated that identification of NSVs that are the causal point of trait variation have been given precedence as a significant area of research.

SPS is the key enzyme controlling the photosynthetic carbon flux towards sucrose (Huber and Huber 1996), and in monocots, five SPS gene families (SPS I to V) have been reported which differ in the number and type of regulatory sites. SPS has been reported to have an N-terminal catalytic glucosyltransferase domain and regulatory phosphorylation sites which have been reported to be associated with light-dark regulation, fructose-6P binding, osmotic regulation and 14-3-3 protein binding (McIntyre et al. 2015).

In the present study, due to complex and highly heterozygous nature of sugarcane genome, a lot of variability was recorded within the coding regions of almost all the genes related to metabolic regulatory networks. The regulatory genes of the sucrose-starch pathway when studied independently between the parents and the bulks, exhibited a significant differential expression for different gene families of SPS, which might be able to explain the regulation of sucrose accumulation machinery in sugarcane. SPS III proteins lack the regulatory phosphorylation sites associated with 14-3-3 protein binding and osmotic stress activation. In present study the eight-fold upregulation of SPS III in the high sucrose parent suggests that SPS III contributes as a housekeeping-like sucrose production gene, which if present in abundance, helps in the overall increase on sucrose till maturity without any interference of external stimulus mainly due to the absence of regulatory site. Similarly, in high sucrose parent, SPS V which is prone to regulation via the 14-3-3 protein binding site, was downregulated two and a half folds. Therefore, it might be possible that in long run, a higher proportion of SPS III coupled with a lower proportion of SPS V enables the sugarcane plants to accumulate higher amounts of sucrose. In majority of the sugarcane varieties it has been observed that the high sucrose accumulation at early stage does not always result in a high sucrose accumulation at the later stage rather plants starts showing maturity symptoms and at the most, a few exceptional varieties may maintain the high sugar content for an extended period. In present study as well the early sucrose bulk didn’t end up in to the highest sucrose bulk, wherein, the SPS III was ten-fold downregulated compared to that of late sucrose bulk. This indicates that this downregulation of SPS III might have diverted the sucrose producing flux towards the other SPS gene family proteins.

In the case of genes associated with sucrose-starch metabolism, 3 small effect NSVs were detected within the glycosyl transferase domain of SPS (Fig. 5), but, no NSV was detected in S6PP-like domain of SPS which has a major role in SPS-SPP binding. On the other hand, in SPP one large effect NSV located in the S6PP C-terminal domain (S6PP-C domain) was identified with a significantly high relative effect (f-value = 0.30). SPP irreversibly catalyses the last step in sucrose synthesis following the formation of sucrose-6-phosphate via SPS. Albi et al. (2016) reported that in higher plants S6PP-C domain is responsible for SPS-SPP dimerization which enhances the efficiency of SPP activity. Hence, it is suggested that the positive effect NSV located in SPP might play an important regulatory role in sucrose metabolism. Similarly, there were NSVs in the other sucrose metabolism genes, *viz*., SuSy, hexokinase, fructokinase which could have a direct effect on the sucrose metabolism pathway.

In C4 plants like sugarcane, phosphoenolpyruvate (PEP) is carboxylated in presence of bicarbonate (HCO-3) to form four carbon acid oxaloacetate (OAA) for an increased efficiency of carbon fixation. Alternatively, PEP can be sequentially metabolized in to pyruvate and acetyl-CoA to produce fatty acids by plastidic pyruvate kinase (PK) inside the chloroplast. (Flugge et al., 2011). The downregulation of plastidic PK in case of high sucrose condition observed in present study could possibly be inhibiting the flux towards fatty acid biosynthesis, and diverting the PEP flux towards an increased degree of carbon fixation.

The overview of regulation pathway revealed upregulation of thioredoxin (TRX), which are small ubiquitous oxidoreductases and plants have an atypical NADP_thioredoxin reductase C (NTRC) localized in plastids containing both NTR and a TRX domain in a single polypeptide. Recently, TRX has been reported to function in the dynamic acclimation of photosynthesis in fluctuating light by promoting light activation of malate dehydrogenase and thus enabling excess reducing power to be exported from chloroplast (Thormahlen et al., 2017). Hence the upregulation of thioredoxin might play a role in improving the source-sink communication and transport resulting higher sucrose accumulating potential.

The SPS genes have diverse expression patterns that are sometimes responsive to environmental stresses, especially cold (Strand et al. 2003) and drought (Yang et al. 2001). The SPS is a major rate limiting enzyme in sucrose synthesis, which is a multi-gene family with variable number of regulatory sites. The present study revealed an upregulation of the cold stress response genes in high sucrose accumulation conditions. This is in conformity with the report of Strand et al. (2003) which stated that cold acclimatization is associated with the tendency of a plant to accumulate higher levels of sucrose. At least two kinases have been implicated in phosphorylation and inactivation of SPS; a Suc non-fermenting 1-related protein kinase (SnRK1), and a calmodulin-like domain protein kinase (CDPK; Winter and Huber, 2000). SPS I, II and V have the phosphorylation site that is inactivated by protein kinase signal during cold stress. In the case of cold stress tolerant genotypes, this inactivation might be reduced in presence of cold stress proteins, which could be a possible reason for the upregulation of cold stress genes in high sucrose producing conditions. In present study, the PS II and photorespiration was downregulated during early sucrose accumulation. McCormick et al. (2006) reported that photosynthetic activity declines during culm maturation in high sucrose accumulating sugarcane cultivars, and suggested that this source-sink communication might be a major regulator of sucrose accumulation. PS II is a stress sensor, and decrease in its activity is the first effect in a plant under stress that induces regulatory process for photochemical quenching, formation of proton gradient, state transitions, downregulation by photoinhibition and gene expression. Matsubara et al. (2002) reported that downregulation of PS II resulted in the activation of cold acclimation in mistletoes, which might also end up as a trigger for higher sucrose accumulation. Efficiency of transport of metabolites is one of the key factors regulating sucrose accumulation. In this study also, the highest degree of upregulation was observed in the genes involved in sugar transport. In this study, transcripts associated with transporters of sugars, peptides and oligopeptides, ABC transporters, SCAMP and mitochondrial solutes were highly upregulated, and similar results have been reported by Matsubara et al. 2002 and Thirugnasambandam et al., 2017. Interestingly, in present study, a negative effect stop codon on the active site of the gene coding for aquaporin was identified. In the case of sugarcane, the level aquaporin transcription has been reported to be variable and genotype-specific (da Silva et al., 2013). Aquaporins are water channel proteins that increase the plant tolerance to stress. Papini-Terzi et al. (2009) also reported a strong association of aquaporins to a high degree of sucrose accumulation. In present study, the low sucrose conditions exhibited an enrichment of aquaporin transcripts containing a premature stop codon located within its amphipathic channel domain (Table 4). This NSV might be playing a major role in reducing the effective amount of aquaporin proteins in sugarcane which might be the cause of reduced sucrose accumulation.

In present study, rather than trying to unravel and follow the complex segregation ratios of NSVs, in the first step, condition-specific differential NSVs were identified that were mapped to the *de novo* assembly. In the case of rice, NSVs have been reported to be more efficient than SNPs from coding as well as non-coding regions, for identifying functional loci and alleles through GWAS (Zhao et al. 2018), wherein, applying such an approach was able to reduce the computational burden and identify functional loci without confounding effect of non-significant loci which were not likely to have a biological function. In the present study, adopting this novel approach of bulk segregant analysis, the background noise in the complex polyploid genomes of sugarcane was minimised, and only the causal elements of trait variation were followed. Although, unlike, diploids, the allele transmission cannot be explicitly modelled for polyploid crops, but results reported here are able to identify initial candidate variations at the genic as well as allelic level, that are responsible for a finer regulation of sucrose accumulation. In a previous study on sugarcane, a similar experimental model exploiting extreme bulks from field grown F_1_ progeny segregating for commercial cane sugar per cent was outlined by Casu et al. (2005). Based on the large number of differentially enriched NSVs identified in this study that were present within the coding regions of important metabolic genes, it is suggested that regulatory variation is predominantly at the allelic level rather than at genic level in the complex polyploid heterozygous sugarcane genome. Moreover, since the bulks used in this study were obtained from a biparental mapping population, a clear picture of the allelic variations was obtained owing to the fact that individuals of the bulk share same pedigree. The availability of transcript information enables association analysis of genes with such important traits in sugarcane as well as their comparative assessment with those from other taxa. Being functionally enriched, the some of NSVs linked to early sucrose accumulation could be the actual cause of variations, and thus could reveal valuable information about gene function leading to a better understanding about causal variations of sucrose accumulation in sugarcane.

## CONCLUSION

In a polyaneuploid genome like sugarcane, where it is not feasible to create a bi-parental structured linkage mapping population (F_2_, RILs and NILs), a novel approach was adopted to investigate BSR-seq data, through which 9905 condition-specific NSVs were identified, of them 43 recorded a high degree of differential enrichment. One of the key findings of this study is that the sucrose phosphate synthase III might play a predominant role in increasing the sucrose accumulation capacity of this crop. The variation at single nucleotide level identified in this study indicates that there are numerous regulatory small effect single nucleotide variations in sugarcane genome, and the trait expression is due to the composite effect of such variations. Sugarcane is not a crop where large effect QTLs could be used for marker-assisted selection, and thus it is suggested to use a cumulative variation-based approach towards genetic gain estimations. Hence, breeding programs based on the composite effect of all non-synonymous genetic variants, located in the coding regions of genes, and their incorporation in selection cycles are expected to fine tune the breeding efficiency for sucrose accumulation.

## Data availability

The raw datasets generated in the present study are available in the NCBI SRA database with accession numbers IISR1101 to IISR1104 under the BioProject PRJNA528576.

## Acknowledgement

This study was conducted with the financial help in the form of research projects from BioCARe, Department of Biotechnology, New Delhi, Government of India to NB, Department of Science and Technology, New Delhi to NB (WOS-A) and SK (SERB). Authors are thankful to the Directors of ICAR-Indian Institute of Sugarcane Research, Lucknow for providing infrastructure facilities, and ICAR-Sugarcane Breeding Institute, Coimbatore, India for allowing to use National Hybridization Garden Facility to execute crossing program. Authors are thankful to Vincent Lacroix, Associate Professor, University of Lyon, France for helpful discussions regarding the data analysis software packages used in the study. Authors are also thankful to The Director, ICAR-National Bureau of Fish Genetic Resources, Lucknow for allowing to use High Performance Computing (HPC) Facility.

## Author contributions

NB, SK conceived and designed the project. NB, AA, SK, PKS, JS, RKS, SK conducted the crossing and plant material development. NB, SK, RKS conducted the experiments. NB, SK performed the sequence data analysis. NB, SK, AA prepared the manuscript. SK coordinated the project.

## Ethical approval

This article does not contain any studies with human participants or animals performed by any of the authors.

## Supplementary Materials

Table S1. Details of the male parent (CoV 92102) specific primer used to screen the true F_1_ individuals from the population.

Table S2. Mean values of sucrose content and other important agronomic traits of 187 F_1_ individuals (data pooled over two years, 2015-16 and 2016-17).

Table S3. List of Differentially expressed genes (DEGs) that were common between the two parents and two bulks along with their log fold change (log FC), log counts per million (log CPM), p-value, structural annotation (Interpro) and assignment to functional bins (Mercator).

Table S4. List of differentially enriched 27626 NSVs identified in this study along with the structural and functional annotation of the contigs hosting these NSVs.

Fig. S1. Annotation of the contigs of the *de novo* assembly using Mercator.

